# Development of infants’ neural speech processing and its relation to later language skills: an MEG study

**DOI:** 10.1101/2021.09.16.460534

**Authors:** T. Christina Zhao, Patricia K. Kuhl

## Abstract

The ‘sensitive period’ for phonetic learning (∼6-12 months) is one of the earliest milestones in language acquisition where infants start to become specialized in processing speech sounds in their native language. In the last decade, advancements in neuroimaging technologies for infants are starting to shed light on the underlying neural mechanisms supporting this important learning period. The current study reports on the largest longitudinal dataset to date with the aim to replicate and extend on two important questions: 1) what are the developmental changes during the ‘sensitive period’ for native and nonnative speech processing? 2) how does native and nonnative speech processing in infants predict later language outcomes? Fifty-four infants were recruited at 7 months of age and their neural processing of speech was measured using Magnetoencephalography (MEG). Specifically, the neural sensitivity to a native and a nonnative speech contrast was indexed by the mismatch response (MMR). They repeated the measurement again at 11 months of age and their language development was further tracked from 12 months to 30 months of age using the MacArthur-Bates Communicative Development Inventory (CDI). Using an *a prior* Region-of-Interest (ROI) approach, we observed significant increases for the Native MMR in the left inferior frontal region (IF) and superior temporal region (ST) from 7 to 11 months, but not for the Nonnative MMR. Complementary whole brain comparison revealed more widespread developmental changes for both contrasts. However, only individual differences in the left IF and ST for the Nonnative MMR at 11 months of age were significant predictors of individual vocabulary growth up to 30 months of age. An exploratory machine-learning based analysis further revealed that whole brain MMR for both Native and Nonnative contrasts can robustly predict later outcomes, but with very different underlying spatial-temporal patterns. The current study extends our current knowledge and suggests that native and nonnative speech processing may follow different developmental trajectories and utilize different mechanisms that are relevant for later language skills.

## 1. Introduction

The *sensitive period* for phonetic learning, roughly between 6 to 12 months of age, is one of the earliest milestones in language acquisition. Learning during this period is a reliable predictor of individual differences in speech and language skills later in life (Kuhl et al., 2008; Kuhl, Conboy, Padden, Nelson, & Pruitt, 2005; Tsao, Liu, & Kuhl, 2004; Zhao, Boorom, Kuhl, & Gordon, 2021). A striking characteristic of this period is the reduction in the ability to discriminate nonnative speech sounds in addition to the improvement in the infants’ ability to discriminate speech sounds in their native language (Kuhl et al., 2006; Werker & Tees, 1984a). Initially, Werker and Tees (1984) demonstrated significantly lower percent of English learning infants were able to discriminate nonnative Hindi and Salish consonant contrasts at 12 months, compared to 6 months of age, while these contrasts remained highly discriminable for almost all Hindi and Salish learning infants. Additional studies revealed a significant increase in native-language speech discrimination performance during the sensitive period, suggesting a learning process (Kuhl et al., 2006; Tsao, Liu, & Kuhl, 2006). Over the decades, similar phenomena have been observed behaviorally in other consonant contrasts, vowels as well as lexical tones (Best & McRoberts, 2003; Kuhl et al., 2006; Kuhl, Williams, Lacerda, Stevens, & Lindblom, 1992; Mattock & Burnham, 2006; Mattock, Molnar, Polka, & Burnham, 2008).

### 1.1 Development of MMR between 6-12 months of age

Despite the ubiquity of the perceptual narrowing phenomenon, the neural mechanisms underpinning perceptual narrowing in infants are largely unresolved and remain a topic of intense research interest (Kuhl, 2010). Over the last decade, the advancement of non-invasive electrophysiological and neuroimaging methods for young children are starting to allow the examination of underlying neural mechanisms. Cheour and colleagues first examined neural discrimination of native vs. nonnative vowel contrasts cross-linguistically in Finnish and Estonian infants at 6 and 12 months of age, using electroencephalography (EEG). They found higher neural sensitivity, as indexed by the mismatch negativity (MMN), to native compared to nonnative vowel contrasts at 12 months but not at 6 months (Cheour et al., 1998). Similar results were later observed in a longitudinal sample with a native and a nonnative stop consonant contrast (Rivera-Gaxiola, Silva-Pereyra, & Kuhl, 2005). That is, the MMN to the native contrast improved, but not nonnative MMN. Both studies reported the MMN effect to be around 250ms-500ms after the speech onset, giving initial clues to the time course of the underlying neural processing of speech contrasts.

In more recent years, the Magnetoencephalography (MEG) has become a more attractive measurement tool, given its advantage over EEG that it not only provides excellent temporal resolution but also good spatial resolution. MEG technology started to allow a closer examination of the underlying neural sources of the mismatch response (MMR). Indeed, the first study with MEG examined infants’ neural responses to speech and nonspeech sounds in newborns, 6-month-olds and 12-month-olds cross-sectionally (Imada et al., 2006). The study provided initial evidence showing the enhancement of neural activation in response to speech, not only in the auditory region (superior temporal) but also in the frontal region (inferior frontal) with increasing age. A second study reported on two separate small cross-sectional datasets comparing 6- to 7-month-olds and 11- to 12-month-olds on two different types of speech contrasts. Both datasets suggest that at the later age, an enhanced MMR was observed in the inferior frontal region for the nonnative contrast when compared to the native contrast, interpreted as an increased difficulty in ‘synthesizing the motor movement’ for speech production (Kuhl, Ramirez, Bosseler, Lin, & Imada, 2014).

### 1.2 Prediction of later language outcomes using MMR

One key prediction regarding the divergence of sensitivity to native vs. nonnative speech contrast is that it sets the stage for later development through native language neural commitment (NLNC) (Kuhl et al., 2008). So far, several pieces of evidence have shown support for this theory. In two complementary studies, Kuhl and colleagues measured 7.5-month-olds’ discrimination to a native and a nonnative speech contrast, first behaviorally (Kuhl et al., 2005) and then using MMN measured with EEG (Kuhl et al., 2008). Critically, a single age in the middle of the ‘sensitive period’ (i.e., 7.5 month) was selected for speech discrimination measurement as it captures the maximum variability across individuals. Then infants’ language development was followed up by the MacArthur-Bates Communication Development Inventory up to 30 months (Fenson et al., 2007; Fenson et al., 1993). The results demonstrated, in both studies, that both native and nonnative speech discrimination at 7.5 months of age predicted later language skills, such as vocabulary and mean length of utterance (MLU), but in *opposite* directions. Particularly, while discrimination of the native contrast was positively correlated with later language outcomes (i.e., better discrimination in infants was linked to faster language growth over time), discrimination of nonnative contrast revealed a negative correlation with later language outcomes (i.e., better discrimination in infants was linked to slower language growth over time). This pattern of results was interpreted to indicate that better nonnative contrast discrimination reflected worse ‘neural commitment’ to the native language and thus indexed a slower native language growth.

One additional EEG study measured infants’ MMN to a native and a nonnative speech contrast at 11 months of age. Their results suggested that while native MMN is not a predictor of later language outcomes, infants with a more mature nonnative MMN demonstrated significantly lower vocabulary scores between 18 and 30 months of age (Rivera-Gaxiola, Klarman, Garcia-Sierra, & Kuhl, 2005).

Thus, these earlier studies suggest that good performance on nonnative contrasts may serve as an indicator of slower later language growth. Most recently, this result was replicated and extended to 6 years of age (Zhao et al., 2021). Infants’ MMR to a nonnative speech contrast was measured in MEG at 11 months of age and infants were followed up at 6 years of age with a comprehensive standardized speech and language assessment by a speech-language pathologist. Infants’ nonnative MMR negatively predicted individual grammar skills as well as their risk of a speech-language disorder at 6 years of age, in line with the NLNC predictions. Native MMR was not measured at 11 months of age in this study.

### 1.3 Current study

Given these two important research questions regarding the ‘sensitive period’ for infant phonetic learning (i.e., the development of native vs. nonnative contrast discrimination and whether performance on either contrast in infancy predicts later language development), no existing dataset allows a truly comprehensive examination of both questions in a well-powered manner. Datasets with multiple measurements of infant discrimination are small (N<15), and largely cross-sectional in nature while datasets with later language follow-ups have only a single measurement of speech discrimination in infancy. The current study reports on the largest dataset to date (N = 54) with infants’ neural discrimination of native and nonnative contrasts measured longitudinally at 7 and 11 months of age using Magnetoencephalography (MEG). Children’s language skills were then followed with CDI from 12-30 months of age. This single dataset allowed us to replicate and extend previous studies and examined two targeted research questions: 1) how speech discrimination of native vs. nonnative contrasts develop from 7- to 11-months of age and 2) whether and how does native and nonnative discrimination performance in infancy predict later language skills.

## Methods

### 2.1 Participants

Fifty-four typically developing infants from monolingual English-speaking households were recruited at 24 weeks to participate in the longitudinal study. The inclusion criteria included the following: 1) full term and born within 14 days of the due date, 2) no known health problems and no more than 3 ear infections, 3) birth weight ranging from 6 to 10 lb., 4) no significant foreign language exposure (i.e., parents and regular caregivers speak only English to the infant) and 5) no previous or concurrent enrollment in infant music classes. All experimental procedures were approved by the University of Washington Institute Review Board and all participating families gave informed consent and were compensated monetarily for their time and effort.

At 7 months of age, all 54 infants (N= 28 male, Mean Age = 201, SD= 8 days) were successful in completing the MEG measurement. However, 4 out of the 54 recordings failed due to technical errors, rendering 50 good quality MEG recordings. At 11 months of age, 48 out of the 54 infants returned for MEG measurement. Out of the 48 MEG measurement sessions (N= 23 male, Mean Age = 338, SD = 7 days), 44 were successful in completion (4 became too fussy to record) and 39 rendered good quality data (5 were excluded due to technical error or excessive noise).

Subsequently, the parents of all 54 infants were followed up at 12, 15, 18, 21, 24, 27 and 30 months to fill out the MacArthur-Bates Communication Development Inventory (Fenson et al., 1993). The descriptive statistics of the CDI info is listed in Table 1.

**Table 1.**
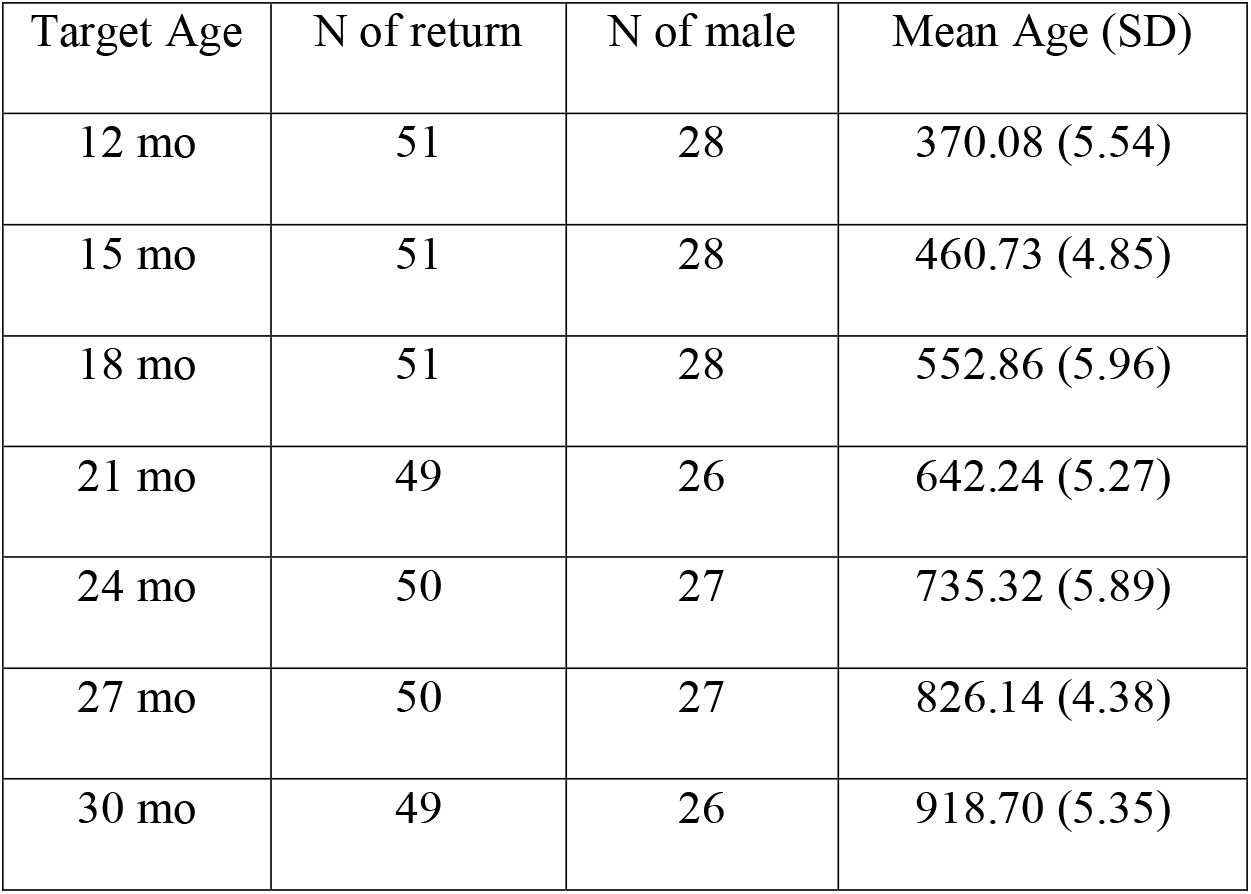
Descriptive statistics of participants for CDI measures at each age.

| Target Age | N of return | N of male | Mean Age (SD) |
| --- | --- | --- | --- |
| 12 mo | 51 | 28 | 370.08 (5.54) |
| 15 mo | 51 | 28 | 460.73 (4.85) |
| 18 mo | 51 | 28 | 552.86 (5.96) |
| 21 mo | 49 | 26 | 642.24 (5.27) |
| 24 mo | 50 | 27 | 735.32 (5.89) |
| 27 mo | 50 | 27 | 826.14 (4.38) |
| 30 mo | 49 | 26 | 918.70 (5.35) |

### 2.2 MEG measurement stimulus

Bilabial stop consonants with varying VOTs were synthesized by the Klatt synthesizer in Praat software (Boersma & Weenink, 2009). The syllable with 0ms VOT was first synthesized with a 2ms noise burst and the vowel /a/. The duration of the syllable is 90ms. The fundamental frequency of the vowel /a/ began at 95Hz and ended at 90Hz. Silent gaps or pre-voicing were added after the initial noise burst to create syllables with +10ms, +40ms and −40ms VOTs, with total durations of 100ms, 130ms and 130ms respectively (Fig 1). The fundamental frequency for the pre-voicing portion was 100Hz. Perception of the synthesized syllables was validated on adult speakers in a previous study using identification and discrimination tasks (Zhao & Kuhl, 2018).

**Figure 1.**
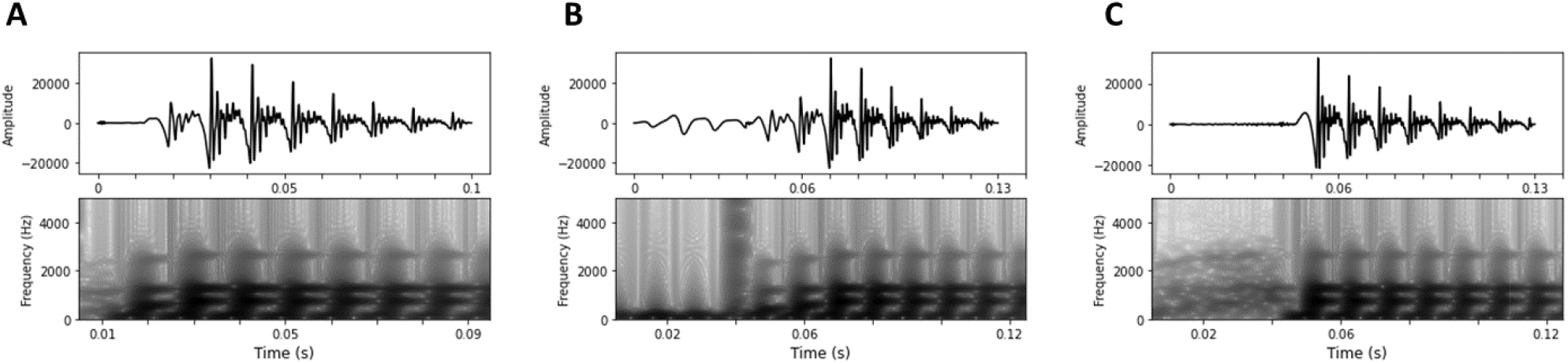
Waveforms (top) and Spectrograms (Bottom) for Standard (/ba/ with +10ms VOT) (A), Nonnative Deviant (/mba/ with - 40ms VOT) (B) and Native Deviant (/pa/ with +40ms VOT) (C).

### 2.3 MEG measurement procedures

Infants were prepared for MEG recordings outside of the magnetically shielded room (MSR) (IMEDCO America Ltd, IN), while a research assistant entertained them. Five head-position-indicator (HPI) coils were attached to the infant’s head that were used to identify head positions under the MEG Dewar. Then, three landmarks (LPA, RPA and Nasion) and the HPI coils were digitized along with 100 additional points along the head surface (Isotrak data) with an electromagnetic 3D digitizer (Fastrak®, Polhemus, Vermont, U.S.A). In addition, a pair of electrocardiography sensors (ECG) were placed on the front and backside of the participants’ left shoulder to record cardiac activity. Infants were then moved and placed in an infant seat made for use in the MEG scanner inside the MSR. The infant’s head was centered and positioned as high as possible relative to the MEG dewar, using foam cushions and padding. The MEG recordings were made with a whole-scalp system with 204 planar gradiometers and 102 magnetometers (VectorView™, Elekta Neuromag Oy, Helsinki, Finland). MEG data was sampled at 1 kHz. During MEG recordings signals from the HPI coils were used to continuously track the child’s head position relative to the MEG sensors.

The sound stimuli were delivered from a TDT RP 2.7 device (Tucker-Davis Technologies Inc., FL), controlled by custom python software on a HP workstation, to a speaker with a flat frequency response. The stimulus was processed such that the RMS values were referenced to 0.01 and it was further resampled to 24,414 Hz for the TDT. The sounds were played at an intensity level of 75 dB measured under the Dewar of the MEG. A multi-feature oddball paradigm was used for stimulus presentation. The syllable with +10ms VOT was used as the Standard (700 trials), the syllables with +40ms and −40ms VOT were used as deviants (150 trials per deviant) with at least two Standards in between Deviants (see Fig 1). The +10ms VOT/+40ms VOT stimulus contrast represents a native phonemic contrast between /ba/ and /pa/, therefore the stimulus with +40ms VOT is from here on referred to as the Native Deviant. On the other hand, the +10ms VOT/-40ms VOT stimulus contrast is a phonemic contrast in nonnative languages, such as Spanish, but not in English. Therefore, the stimulus with −40ms VOT is from here on referred to as the Nonnative Deviant. The stimulus-onset-asynchrony (SOA) was 600ms with jitters within a 100ms window. The whole recording took around 10 minutes of time during which the participants listened passively while a research assistant used silent toys to keep the infants calm and attentive to the toys (Fig 2A).

**Figure 2.**
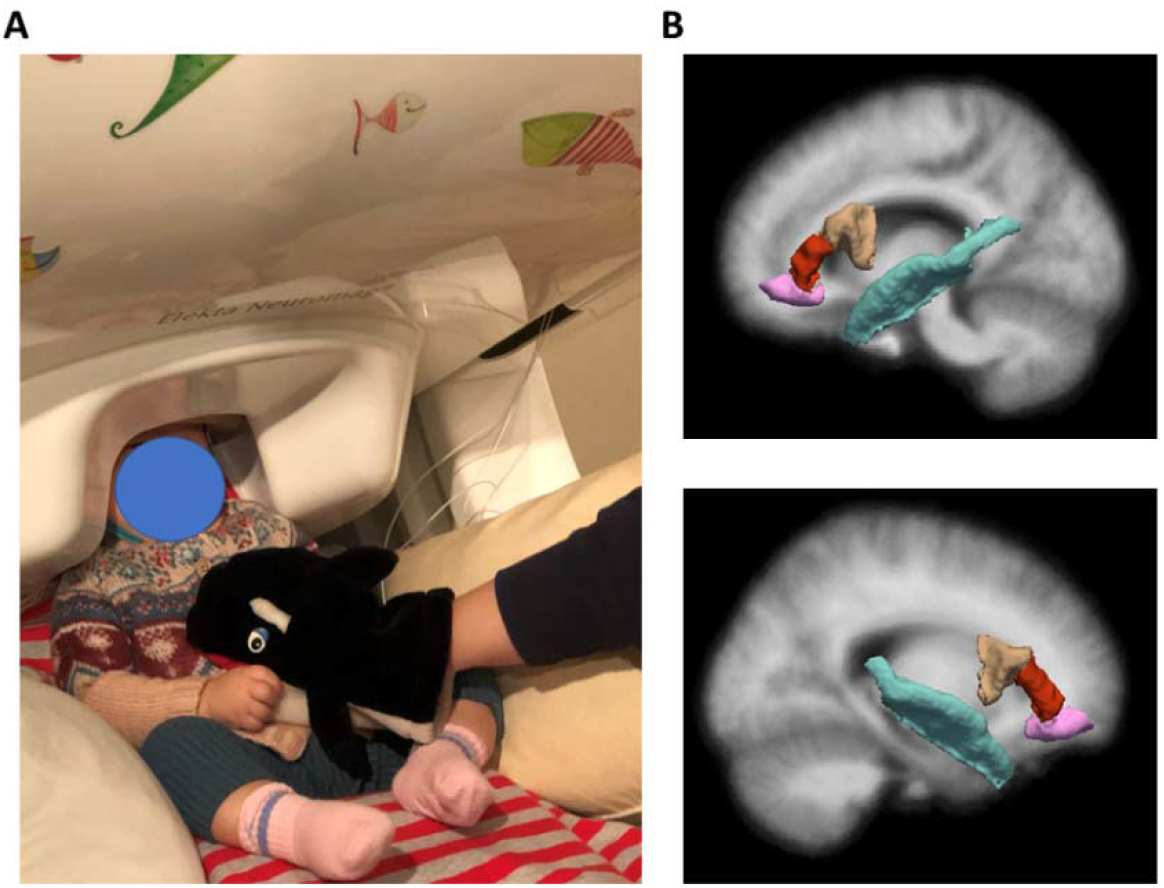
(A) A 7-month-old infant being prepared to start MEG recording. (B) The ROIs selected a priori for analyses.

### 2.4 CDI

Productive vocabulary was assessed when participants reached the age of 12 months and continued to 30 months of age using the MacArthur-Bates Communicative Development Inventory (CDI) at 3-month intervals. For 12 and 15 months, the ‘Words and Gesture’ version was used and starting at 18 months, the ‘Words and Sentences’ version was used (Fenson et al., 2007; Fenson et al., 1993). Parents were given a 4-week window around the target dates to respond, and the averaged ages of the surveys are listed in Table 1. Two specific measures were taken: 1) Vocabulary: number of words produced from 12 to 30 months and 2) M3L (mean of the 3 longest utterances) from 18 and 30 months. Specifically, vocabulary was measured by parents’ checking all the words the children can produce on a word list on the survey, with a maximum number of 680 words. For M3L, parents were instructed to write down the children’ longest 3 utterances and the average length of the utterances was calculated.

### 2.5 Data Analyses

#### 2.5.1 MEG data processing

All MEG data processing was done using the MNE-python software v0.23 (Gramfort et al., 2014). The raw MEG recordings underwent a series of standardized preprocessing steps for noise suppression. The temporal signal space separation (tSSS) and head movement compensation were used first to suppress noise from outside of the MEG dewar and to compensate for effects related to infants’ head movement during the recording (Taulu & Hari, 2009; Taulu & Kajola, 2005). This procedure is designed to improve the signal-to-noise ratio of the data by suppressing external interference (i.e., noise from outside of the helmet) without introducing excessive reconstruction noise. Then, the signal-space projection (SSP) method was adopted to isolate components of physiological artifacts (i.e., heartbeats) (Uusitalo & Ilmoniemi, 1997). Lastly, the signal was low-pass filtered at 50 Hz with dead channels rejected.

Epochs were identified for all types of stimuli and were selected from 100ms before stimulus onset to 600ms after stimulus onset. For both deviants, evoked responses were calculated by averaging across all epochs after rejecting any epochs with peak-to-peak amplitude over 4 pT/cm for gradiometers or 4 pT for magnetometers. For standards, only epochs immediately before a deviant were first selected and then 150 epochs were further randomly selected to calculate the evoked responses following the same procedure. This selection ensured the evoked responses for Standards and the two Deviants have similar signal-to-noise ratios.

To estimate the location of neural generators underlying the evoked responses, each subject’s anatomical landmarks and additional scalp points were used with an iterative nearest-point algorithm to rescale a surrogate age-appropriate template MRI to match the subject’s head shape (O’Reilly, Larson, Richards, & Elsabbagh, 2021). FreeSurfer was used to extract the inner skull surface (watershed algorithm) and the cortical and subcortical structures segmented from the surrogate MRI (Dale, Fischl, & Sereno, 1999). A one-layer conductor model based on the rescaled inner skull surface was constructed for forward modeling (Hämäläinen & Sarvas, 1989). For 7-month-old recordings, a volumetric source space consisted of 5981 dipoles evenly spatially distributed based on a 5mm grid within the inner skull surface of the 6-month-old template, while for 11-month-old recordings, a volumetric source space consisted of 7417 dipoles were used (O’Reilly et al., 2021). Because surrogate head models and source spaces were used for each subject, source orientations were unconstrained (free orientation). Baseline noise covariance was estimated using empty room recordings made on the same day of the MEG session. Dipolar currents were estimated from the MEG sensor data using an anatomically constrained minimum-norm linear estimation approach to obtain dSPM values at each source location (Dale et al., 2000). Two mismatch responses (MMRs) were subsequently calculated at the source level by subtracting the Standards from each Deviant. That is, a Native MMR was calculated by subtracting the vectors of Standard from the vectors of Native Deviant and then the magnitude of the vectors was calculated. Similarly, a Nonnative MMR was calculated using Standard and Nonnative Deviant.

To aggregate group level data for each age, each participant’s MMRs were further morphed from their individual source space to the fsaverage template within the volumetric source space. The Desikan-Dilliany Atlas (i.e. ‘aparc’ atlas in Freesurfer) was then applied to reduce the data into 114 labels by averaging across vertices within each label (Desikan et al., 2006). Four regions-of-interest (ROIs) were *a priorly* identified based on existing literature for further statistical analysis: left and right inferior frontal region (i.e., pars orbitalis, pars opercularis, pars triangularis labels combined) and superior temporal region (i.e., superior temporal label) (Fig 2B). All data reported in this study are publicly available at Open Science Framework (Zhao, 2021).

## 3. Results

### 3.1 Developmental change of MMRs

To address the first research question regarding the developmental change of MMRs, we adopted two analyses approaches. 1) The Region-of-Interest (ROI) approach (section 3.1.1), where spatial regions as well as time windows were selected *a prior* based on existing literature for MMR, was first conducted to better compare our current results with the existing literature. 2) Then, the whole-brain approach (section 3.1.2) that is more exploratory in nature, was also conducted to help identify additional regions and time windows that undergo significant changes from 7 to 11 months of age.

#### 3.1.1 The ROI Approach

Based on the existing literature, the Region-of-Interest (ROI) approach examined the MMRs in the 4 spatial areas, namely, left and right auditory region (ST) and left and right frontal region (IF). The time series of the MMRs averaged across these 4 spatial ROIs for the Native and Nonnative contrasts can be visualized in Fig 3A for 7 months of age and Fig 3B for 11 months of age. The time window of 250- 500ms was also selected *a priori* based on previously published papers (Conboy & Kuhl, 2011; Kuhl et al., 2014; Rivera-Gaxiola, Silva-Pereyra, et al., 2005). MMRs averaged across the time window for all the ROIs at each age group can be visualized in Fig 3C.

**Figure 3.**
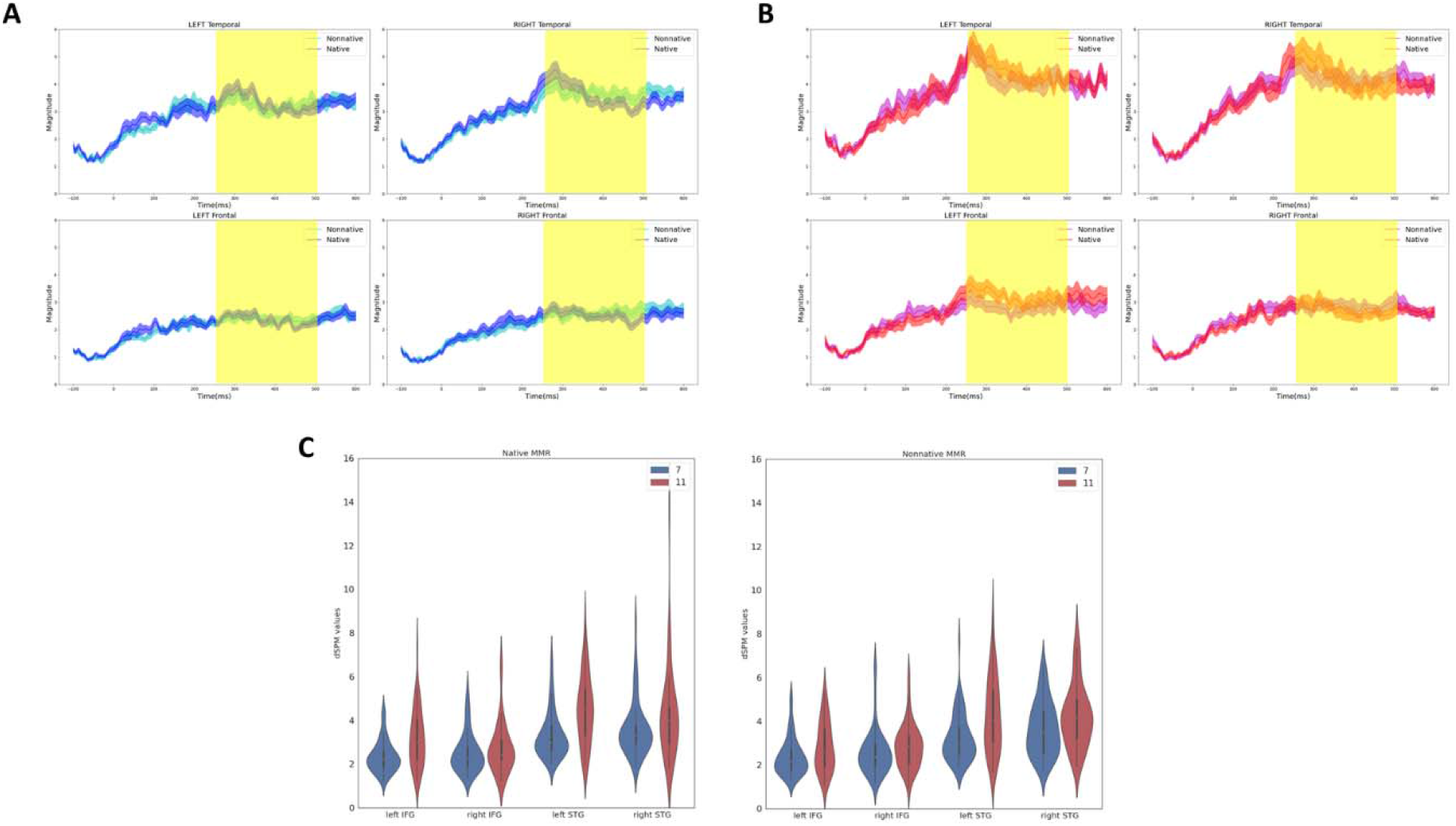
Mismatch responses at 7 (A) and 11 (B) months of age in the four regions-of-interest (ROIs). The yellow shaded region indicates the time window (250-500ms) used for mismatch response. (C) Violin plots of averaged MMR across the time window in ROIs at 7 (blue) and 11 (red) months of age, for the native (left) and nonnative (right) contrasts.

Two types of statistical analysis were used to examine the developmental change in MMRs from 7- to 11-months of age given the missing data at 11 months of age. The first approach utilized the subset of the data (N=35) where each participant had good data at both ages. Paired t-tests were used to test for differences between ages for the MMR in a specific ROI for either the Native or Nonnative contrast. The results are listed in Table 2 (left side). The second approach utilized a linear mixed-effect model with age as the fixed factor and individual participant as a random factor. This approach allows all available data to be utilized (Table 2 right side). Both analyses were done using R software in conjunction with the nlme package (Pinheiro, Bates, DebRoy, Sarkar, & Team, 2021; R Core Team, 2020). The results from both analyses converged and suggested that the MMR significantly increased from 7 to 11 months age in the left IFG and left STG only for the Native contrast. There is a marginal increase in the left STG for the Nonnative contrast given that it was only significant in the linear mixed-effect model.

**Table 2.**
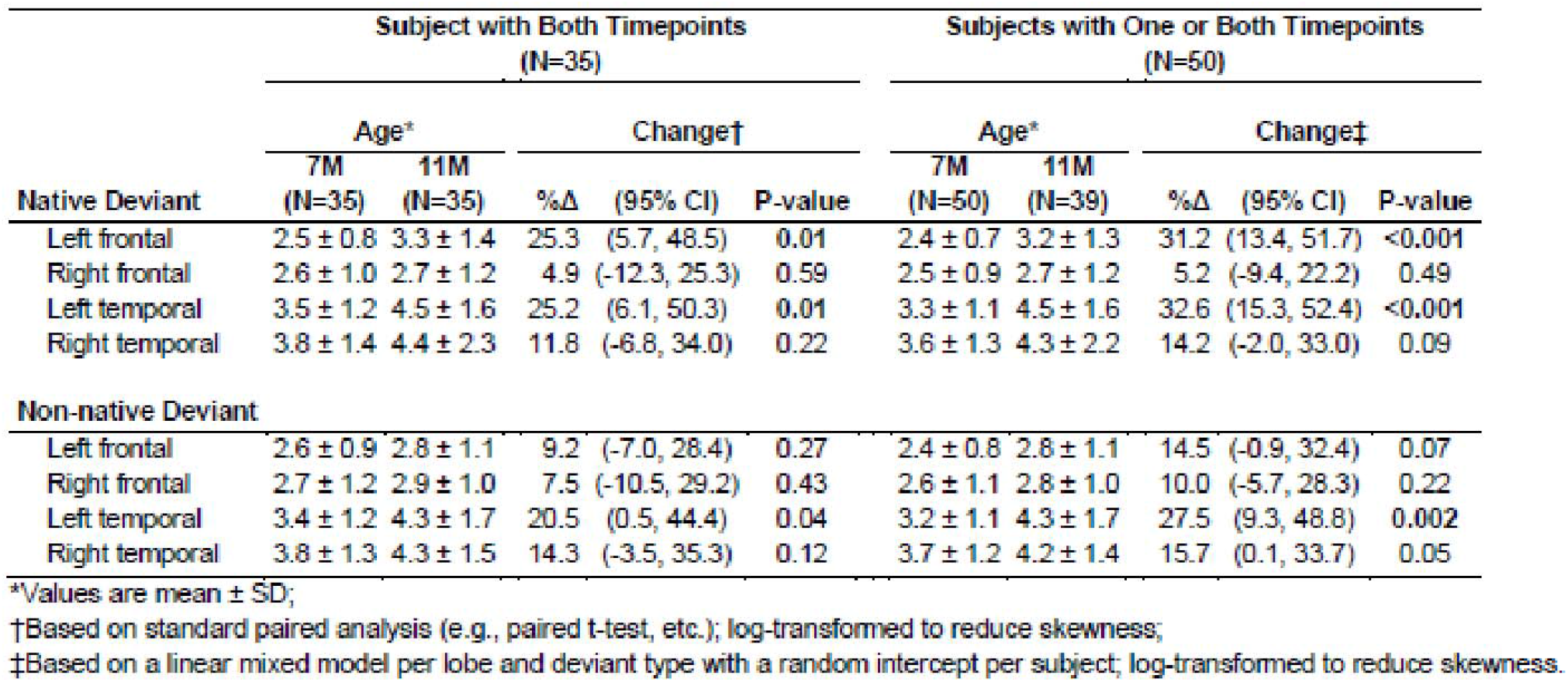
Change in MMR values for each deviant type in ROIs from 7 to 11 months of age. Note while the p values reported are not corrected for multiple comparison, the marking of significance (bold) was done after Bonferroni-correction.

|  | Subject with Both Timepoints<br>(N=35) |  |  |  |  | Subjects with One or Both Timepoints<br>(N=50) |  |  |  |  |
| --- | --- | --- | --- | --- | --- | --- | --- | --- | --- | --- |
|  | Age* |  | Change† |  |  | Age* |  | Change‡ |  |  |
|  | 7M<br>(N=35) | 11M<br>(N=35) | %Δ | (95% CI) | P-value | 7M<br>(N=50) | 11M<br>(N=39) | %Δ | (95% CI) | P-value |
| **Native Deviant** |  |  |  |  |  |  |  |  |  |  |
| Left frontal | 2.5 ± 0.8 | 3.3 ± 1.4 | 25.3 | (5.7, 48.5) | **0.01** | 2.4 ± 0.7 | 3.2 ± 1.3 | 31.2 | (13.4, 51.7) | **<0.001** |
| Right frontal | 2.6 ± 1.0 | 2.7 ± 1.2 | 4.9 | (-12.3, 25.3) | 0.59 | 2.5 ± 0.9 | 2.7 ± 1.2 | 5.2 | (-9.4, 22.2) | 0.49 |
| Left temporal | 3.5 ± 1.2 | 4.5 ± 1.6 | 25.2 | (6.1, 50.3) | **0.01** | 3.3 ± 1.1 | 4.5 ± 1.6 | 32.6 | (15.3, 52.4) | **<0.001** |
| Right temporal | 3.8 ± 1.4 | 4.4 ± 2.3 | 11.8 | (-6.8, 34.0) | 0.22 | 3.6 ± 1.3 | 4.3 ± 2.2 | 14.2 | (-2.0, 33.0) | 0.09 |
| **Non-native Deviant** |  |  |  |  |  |  |  |  |  |  |
| Left frontal | 2.6 ± 0.9 | 2.8 ± 1.1 | 9.2 | (-7.0, 28.4) | 0.27 | 2.4 ± 0.8 | 2.8 ± 1.1 | 14.5 | (-0.9, 32.4) | 0.07 |
| Right frontal | 2.7 ± 1.2 | 2.9 ± 1.0 | 7.5 | (-10.5, 29.2) | 0.43 | 2.6 ± 1.1 | 2.8 ± 1.0 | 10.0 | (-5.7, 28.3) | 0.22 |
| Left temporal | 3.4 ± 1.2 | 4.3 ± 1.7 | 20.5 | (0.5, 44.4) | 0.04 | 3.2 ± 1.1 | 4.3 ± 1.7 | 27.5 | (9.3, 48.8) | **0.002** |
| Right temporal | 3.8 ± 1.3 | 4.3 ± 1.5 | 14.3 | (-3.5, 35.3) | 0.12 | 3.7 ± 1.2 | 4.2 ± 1.4 | 15.7 | (0.1, 33.7) | 0.05 |

\*Values are mean ± SD;
†Based on standard paired analysis (e.g., paired t-test, etc.); log-transformed to reduce skewness;
‡Based on a linear mixed model per lobe and deviant type with a random intercept per subject; log-transformed to reduce skewness.

#### 3.1.2 The whole brain approach

A second, more exploratory approach was also used to examine the developmental change of MMR from 7 months to 11 months of age at the whole brain level for Native and Nonnative contrasts separately. By taking into consideration the spatial-temporal pattern of the MMRs, the analyses may inform any developmental changes that are outside of the *a priori* selected ROIs (i.e., spatial regions and time windows). Specifically, the non-parametric cluster-level paired t-test was conducted with the threshold-free cluster enhancement method (TFCE) (Smith & Nichols, 2009), using the subset of the data where participants have both 7- and 11-month data (N=35).

Significant differences between 7- and 11-month MMRs for both contrasts can be visualized in Fig 4 and in supplemental movies. Critically, peak differences were observed within the 250-500ms with the Native MMR peaking later than the Nonnative MMR, confirming the robustness of the time window selected for the ROI analyses. Overall, at the peak difference, the Native MMR shows much more left lateralized change (Fig 4B) than the Nonnative MMR (Fig 4A). However, the spatial regions involved were more widespread than the ST and IF regions from existing literature, with large changes potentially in regions such as sensorimotor as well as subcortical regions (e.g., brainstem).

**Figure 4.**
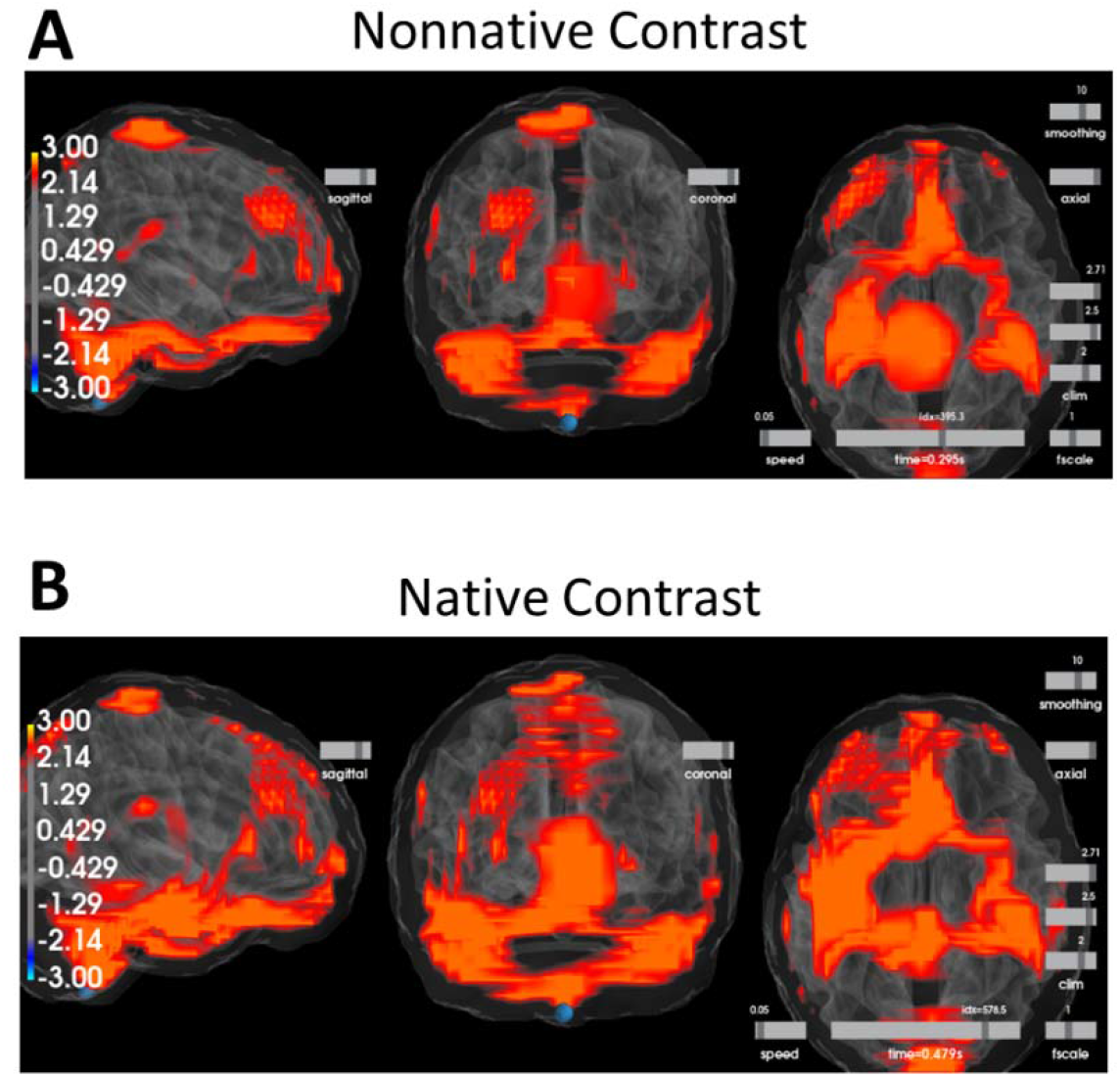
Peak differences between 7- and 11-month MMR at the whole brain level for A) the nonnative contrast and B) the native contrast.

### 3.2 MMRs predicting later language outcomes

To address our second research question, we first adopted an existing approach and examined whether MMRs in each ROI predict the growth of language skills (i.e., vocabulary and M3L). Extending from previous research, we did the same analyses for 7- and 11-month MMR (section 3.2.1). Further, we utilized a new exploratory approach that examines the whole-brain MMR for the entire time series which allows us to identify additional spatial-temporal patterns in MMR that can predict later language outcomes (section 3.2.2).

#### 3.2.1 The ROI approach

The growth of vocabulary was modeled by linear mixed effect models using data from 12 to 30 months of age. Separate tests were conducted to examine whether MMRs in each ROI at each age were predictive of language growth. The vocabulary values were transformed before modeling to account for right-skewness, saturation of the word list, and non-linearity. Specifically, the vocabulary values were first divided by its maximum value to change it to a proportion; the arcsine-transformation, a common variance-stabilizing transformation for proportions, was then applied; lastly, the result was standardized to have 0 mean and unit standard deviation (Bosseler et al., 2021).

Results are listed in Table 3. Only the Nonnative MMRs in the left IF and left ST at 11 months of age significantly predicted the individual growth of vocabulary from 12 to 30 months of age. The results can be further visualized in Fig 5. Using a median split, individuals with larger Nonnative MMRs (green lines) in left IF (Fig 5A) and left ST (Fig 5B) have a larger number of words produced across developmental age. The same trends can be observed for MMRs to the native contrast at 11 months of age but they did not reach significance. No predictive relations were observed between 7-month MMRs and later vocabulary. Similarly, the same approach was applied to M3L between 18 to 30 months of age. The results are listed in supplemental material.

**Table 3.** MMR in ROIs predicting growth of vocabulary from 12 to 30 months of age.

| Native Deviant | Age 7M<br>(N=47) |  |  | Age 11M<br>(N=39) |  |  |
| --- | --- | --- | --- | --- | --- | --- |
|  | B | CI | P-value | B | CI | P-value |
| Left frontal | 0.01 | (-0.07, 0.10) | 0.72 | 0.09 | (-0.00, 0.18) | 0.052 |
| Right frontal | -0.00 | (-0.09, 0.08) | 0.98 | 0.09 | (0.00, 0.18) | 0.050 |
| Left temporal | 0.03 | (-0.05, 0.11) | 0.48 | 0.10 | (0.01, 0.18) | 0.035 |
| Right temporal | -0.02 | (-0.11, 0.06) | 0.56 | 0.06 | (-0.03, 0.15) | 0.20 |
| Nonnative Deviant |  |  |  |  |  |  |
| Left frontal | 0.06 | (-0.03, 0.14) | 0.17 | 0.12 | (0.04, 0.21) | 0.005 |
| Right frontal | 0.01 | (-0.07, 0.10) | 0.74 | 0.06 | (-0.03, 0.16) | 0.17 |
| Left temporal | 0.06 | (-0.02, 0.15) | 0.13 | 0.12 | (0.03, 0.20) | 0.009 |
| Right temporal | 0.02 | (-0.07, 0.10) | 0.67 | 0.03 | (-0.06, 0.13) | 0.50 |

**Figure 5.**
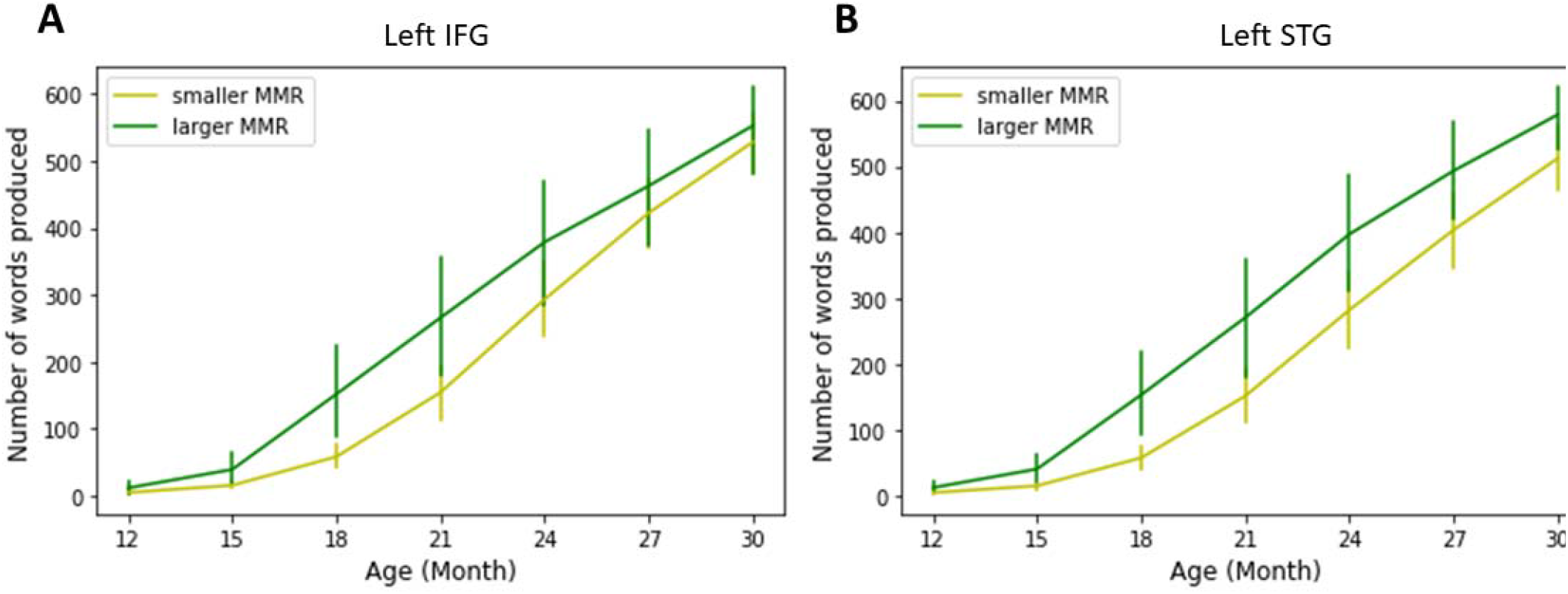
Growth curve of number of words produced from 12 month to 30 months of age, based on a median split of MMR values in left IF (A) and left ST (B) for the nonnative contrast.

#### 3.2.2 The whole brain approach

A second, more exploratory approach was also used to examine the relationship between whole-brain MMR at 11 months of age and vocabulary scores at 24 months of age. By taking into consideration the spatial-temporal pattern of the MMRs, the analyses may inform any predictive patterns that are outside of the *a priori* selected ROIs and time window. The 24-month vocabulary score was selected due to its lack of skewness, floor, or ceiling effects in the data.

Specifically, we employed a machine-learning based method that utilized the whole-brain spatial-temporal pattern of the MMR at 11 months to predict each individual’s vocabulary score at 24 months. The machine-learning analyses were done using the open source scikit-learn package (Pedregosa et al., 2011) in conjunction with the MNE-python software. This method was an extension from a previously published study (Zhao et al., 2021). Specifically, for each time sample, we employed a support-vector regression (SVR) where the model uses MMR values from all 114 label regions, therefore taking spatial pattern of the MMR into consideration, to predict individual 24-month vocabulary scores (Drucker, Burges, Kaufman, Smola, & Vapnik, 1996). The dataset is first randomly split into a training and a testing set. The MMR spatial-temporal patterns in the training set were first used to fit the model with a linear kernel function (C=1.0, epsilon =0.1). Once the model is trained, the MMR spatial-temporal patterns from the testing set were then used to generate predictions of the vocabulary value. A leave-one-out cross-validation method was used to enhance model prediction. The R^2^ coefficient of determination between actual measured Vocabulary and model predicted Vocabulary is taken as an index of model performance. The same process was repeated for every time sample of the MMR, which generates a temporal sequence of R^2^ (Fig 4 BC, upper left column).

To further evaluate the model performance, within each time sample, we shuffled the correspondence between the Vocabulary score and MMR spatial pattern across individuals and then conducted the same SVR analyses. In such cases, the MMR spatial pattern should bear no predictive value to Vocabulary score and the R^2^ should reflect a model performing at chance level. We repeated this process 100 times for each time point. Then we generated an empirical null distribution of R^2^ by pooling all the R^2^ values from each time point (i.e. 100 permutations for each time point) and we compared our originally obtained R^2^ coefficient against this distribution (Xie, Reetzke, & Chandrasekaran, 2019). Specifically, we considered that if our originally obtained R2 value is larger than the 95% percentile of the empirical null distribution (red line in Fig 5) at a specific time point, then the spatial pattern of that time point can significantly predict Vocabulary of an individual. A final SVR was then fit by all the time points that were deemed significant by permutation.

Using this method, we examined 11-month MMR for both Native and Nonnative contrasts and their prediction of 24-month Vocabulary. For the Nonnative MMR, only the time window 324-325ms was found to predict individual Vocabulary significantly (Fig 6A left column, cyan shaded regions). The scatter plot of the model predicted Vocabulary and the actual Vocabulary can be visualized in Fig 6A right column (r = 0.42, p = 0.008, R^2^ = 0.16). The spatial regions with the highest model coefficients (top 10%) can be visualized in Fig 6A bottom panel. For the Native MMR, more time windows (7-13ms, 17- 20ms, 42-47ms, 99-109ms, 152-163ms, 177-178ms) emerged from permutation to predict individual Vocabulary significantly (Fig 6B left column, cyan shaded regions). The scatter plot of the model predicted Vocabulary and actual vocabulary can be visualized in Fig 6B right column (r = 0.53, *p* <0.001, R^2^ = 0.26). The spatial regions with the highest model coefficients (top 10%) can be visualized in Fig 6B bottom panel.

**Figure 6.**
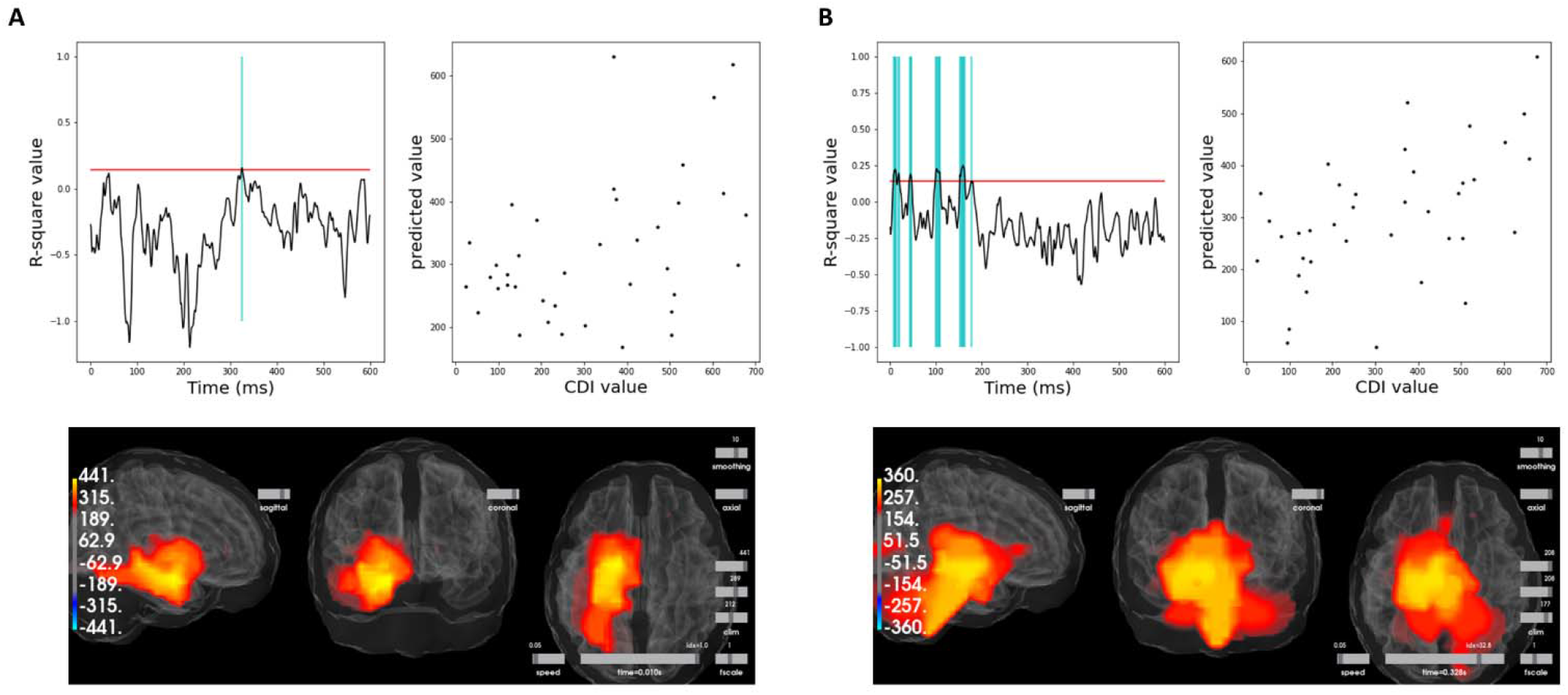
Relations between 24mo vocabulary and whole brain mismatch response for nonnative contrast (A) and native contrast (B).

Overall, the whole-brain MMR for both contrasts at 11 months can generate robust predictions of individual vocabulary values at 24 months. The time window and spatial pattern for the Nonnative MMR are largely in line with the ROI analyses. That is, the left lateralized pattern in the MMR time window is likely to be the driving force of the predictive relation. However, the spatial-temporal pattern for the Native MMR paints a different picture. A much earlier time window (<200ms) in a deeper source (e.g., brainstem) seems to be driving the predictive relation, explaining why no significant relation was observed in the ROI growth curve analyses.

## 4. Discussion

The current study significantly extends our current understanding of the neural development underlying native and nonnative speech discrimination during the ‘sensitive period,’ and more importantly, sheds light on the association between early speech discrimination and later language development. The current study reported on the largest longitudinal dataset to date and comprehensively addressed both questions. Specifically, mismatch responses (MMR) to a native and a nonnative speech contrast were measured using MEG longitudinally at 7- and 11-months of age in monolingual English-learning infants. Later language skills were followed up from 12-to-30 months of age using the CDI. First, by using the same ROI analyses approach as previous studies, the current study demonstrated that only the Native MMR in the left ST and IF significantly increased from 7- to 11-months of age. Nonnative MMR remained largely unchanged with some marginal increase. However, the whole brain comparison revealed much more widespread changes from 7- to 11-month for both contrasts, including additional regions in the sensorimotor as well as subcortical regions. Using the same ROIs, we then examined whether the MMRs predicted the growth of vocabulary and M3L up to 30 months of age. The results show that only the Nonnative MMR in the left ST and IF at 11 months of age were significant predictors of vocabulary growth. Native MMRs at 11 months as well as all MMRs at 7 months of age were non-predictors of vocabulary growth. A similar pattern was also observed for M3L, with the predictive strength between Nonnative MMR restricted to left frontal and the effect marginal (see supplemental material). Additionally, the whole brain approach revealed much earlier time windows in the Native MMR to be highly predictive of later language outcome, suggesting potentially different mechanisms at play.

### 4.1. Development of speech discrimination

The current study reported a robust increase of the native MMR from 7- to 11-months of age that is highly left lateralized. The increase of native MMR is in line with the two previous EEG studies that reported on developmental changes of MMN with longitudinal samples (Cheour et al., 1998; Rivera-Gaxiola, Silva-Pereyra, et al., 2005). Together, the increase in MMR supports the behavior findings that infants’ discrimination of native contrasts improves during the sensitive period (Kuhl et al., 2006; Tsao et al., 2006). Extending from the previous EEG results, the current results demonstrated that the increase of Native MMR is lateralized to the left hemisphere from both the ROI and the whole brain analyses approach. This left lateralization is largely in line with speech perception models (Hickok & Poeppel, 2016).

In contrast, the current study reported on a Nonnative MMR development that is different from previous studies. Specifically, the ROI analyses suggested a largely unchanged Nonnative MMR with marginal increase in the left ST while the whole brain analyses revealed more widespread increases from 7- to 11-months of age. This result is different from behavioral results as well as previous EEG results suggesting a decline of sensitivity/MMN to nonnative contrasts from 6 to 12 months of age (Best & McRoberts, 2003; Cheour et al., 1998; Kuhl et al., 2006; Rivera-Gaxiola, Silva-Pereyra, et al., 2005; Werker & Tees, 1984a). More recently, a decline in early sensory encoding of nonnative speech was also reported from 7- to 11-months of age (Zhao et al., in prep).

This discrepancy may be largely attributed to the fact that our experimental paradigm may have elicited more acoustic processing than phonetic processing, particularly for the Nonnative contrast. The increase in acoustic processing is caused by three factors. 1) While efforts were made to tightly control the time course of the two Deviants, therefore making Native and Nonnative MMR more comparable to each other, both Deviants were 30ms longer than the Standard. This duration difference between Standard and Nonnative deviant may have increased the portion of acoustic processing in the Nonnative MMR, which may undergo enhancement during early development, in comparison to phonetic processing that declines during this period. 2) The different choices of nonnative speech contrasts across different studies may also have elicited differential processing. We will discuss this point more in depth in a later section (4.3.1). 3) The shorter Stimulus-Onset-Asynchrony (SOA) used compared to previous studies may have also driven a more acoustic processing than phonetic processing.

On the other hand, it is also possible that the decline in behavioral discrimination for nonnative contrasts is reflected in the underlying neural processes as a maintenance (i.e., no improvement) in the left IF region. While parallel results have been demonstrated between behavioral and MMR measurement, a direct correlation between MMR and behavioral discrimination is hardly reported. Further, previous studies using EEG measured largely reported effects from central and frontal electrodes where the underlying source can be a combination of auditory and frontal regions. Future research is needed to examine a direct link between behavioral discrimination of speech contrasts and its underlying neural sources to help better understand the development of speech processing both in behavior and in the neural processes.

### 4.2 Prediction of later language outcome

The current study provided a unique opportunity to compare the predictive relations to later language outcome between 7-month and 11-month MMRs. From the ROI analyses, there is clear evidence that 11-month MMR, but not 7-month MMR, can robustly predict language growth up to 30 months. This contrast further bolsters the idea of the ‘sensitive period’ where infants’ neural function undergoes significant changes, and the outcome of this period is what matters in setting infants up on a good language learning trajectory.

Further, a divergence was observed in predictive relations between the Native and Nonnative MMR at 11-month, similar to a previous study (Rivera-Gaxiola, Klarman, et al., 2005). Only Nonnative MMRs in left IF and ST regions were significant predictors while the Native MMRs only trended in the same direction. These differences between Native and Nonnative MMR are likely due to a reduced level of variability in the Native MMR. In the previous studies (Kuhl et al., 2008; Kuhl et al., 2005), a single age (i.e., 7.5 month) was chosen in the middle of the ‘sensitive period’ and both native and nonnative speech discrimination were shown to predict later language outcomes. Compared to 7.5 month, most infants have mastered the native contrast at 11 months of age (i.e., significant enhancement in Native MMR at the group level) and thus the variability across participants may have been significantly reduced for native MMR to be a good predictor of later language growth.

While the Nonnative MMR in the left ST and IF regions at 11 months were significant predictors of later vocabulary growth in the current study, the direction of prediction is different from previous studies. Specifically, the Nonnative MMR predicts later vocabulary in a *positive* direction (i.e., better MMR, faster vocabulary growth) while previous studies demonstrate a *negative* direction (i.e., better MMR, slower vocabulary growth) (Kuhl et al., 2008; Zhao et al., 2021). This difference, again, may be due to the increased acoustic processing for the Nonnative contrast discussed in 4.1. Taken together, the *positive* correlation observed in the current study (i.e., better nonnative MMR, faster language growth) may indicate a mechanism in which better general auditory processing is predictive of later language development, similar to many other previous studies focusing on general auditory processing (Cantiani et al., 2019; Molfese & Molfese, 1997).

### 4.3 Importance of complementary ROI and whole brain analyses

The whole brain approach in the current study allowed examination of the whole spatial-temporal patterns of the brain that are involved in language processing instead of focusing on specific ROIs within a specific time window. While examination of whole spatial patterns does not allow total disambiguation of different neural sources due to point spread in MEG, it provides an important complementary view to ROI analyses with regard to the extent of the effect. Indeed, both whole brain analyses employed in the current study provided new information.

First, the whole brain comparison between 7- and 11-month MMR revealed additional regions that underwent significant changes between ages, especially in sensorimotor regions as well as subcortical regions. The sensorimotor region has been shown to be very relevant for speech processing in infants and the development of its function may support infants as they learn to produce the speech sounds they have been hearing (Choi, Bruderer, & Werker, 2019; Choi, Dehaene-Lambertz, Peña, & Werker, 2021; Kuhl, 2021).

Second, the whole brain MMR machine-learning approach also provided new insights on the relation between infant speech processing and later language outcomes. While no significant correlation was found between Native MMR and later language development using ROI analysis, a different spatial-temporal pattern in the Native MMR time-series was identified to be highly predictive of later vocabulary score. Specifically, the pattern consists of a much earlier temporal window (<200ms) in a much deeper region (i.e., subcortical and brainstem). What this may suggest is that the Native MMR at 11 months differentiates individuals not based on whether they can process native speech sounds phonemically (i.e., based on the later 250-500ms time window), but rather, based on the early sensory encoding of the fine acoustic properties of the speech sounds.

The potentially important role of subcortical processes in predicting later language outcomes parallels the significant developmental change in subcortical regions from 7- to 11- months in the whole brain analyses. Indeed, recent growth in the adult research literature has demonstrated repeatedly that early sensory encoding of speech in largely subcortical regions is highly relevant for speech discrimination and highly malleable through experience (Slabu, Grimm, & Escera, 2012; Zhao & Kuhl, 2018; Zhao, Masapollo, Polka, Ménard, & Kuhl, 2019). This result opens many more questions that future research will need to elucidate. For example, what is the relation between early sensory encoding of speech and later discrimination of speech? Does the trajectory follow similar or different developmental patterns for native and nonnative speech?

### 4.4 Limitation and future directions

#### 4.4.1 Selection of nonnative contrasts

The current study is limited by the single nonnative deviant chosen for measuring the Nonnative MMR. Specifically, this contrast (i.e., voiced vs. pre-voiced stop consonant) represents a phonemic contrast in languages such as Spanish, but not in English. Yet, pre-voicing is still largely present in the production of voiced bilabial stop consonants in English speakers (Fish, García-Sierra, Ramírez-Esparza, & Kuhl, 2017; Lisker & Abramson, 1964). Thus, while this contrast is a more difficult contrast to process as it is not used phonemically in English, English-learning infants may still maintain sensitivity to the acoustic differences as they may be relevant for speaker identification or evaluating the goodness of the token in the category.

This leads to the question of what characterizes a ‘nonnative’ contrast for English speakers/learners. Indeed, a recent all-encompassing review compared infants’ perception of a range of native and nonnative contrasts and showed that discrimination of the contrasts varies depending on the contrast (Best, Goldstein, Nam, & Tyler, 2016). The Mandarin Chinese alveolo-palatal affricate /tchi/ and fricative /ci/ contrast was used as the nonnative contrast in a series of behavioral and M/EEG studies before (Kuhl et al., 2008; Kuhl et al., 2005; Kuhl et al., 2014). Critically, evidence supporting the NLNC hypotheses, whereby the nonnative contrast discrimination *negatively* predicted later language outcome (i.e., better nonnative contrast discrimination, slower language growth), largely used this Mandarin Chinese alveolo-palatal affricate vs. fricative contrast (Kuhl et al., 2008; Kuhl et al., 2005). However, it is worth noting that while English adult speakers shown to have poorer performance in discriminating these contrasts than native Mandarin speakers, similar place of articulation cues are used in English and the English speakers demonstrated fairly high sensitivity to the Mandarin contrast (Tsao et al., 2006).

Our current study along with several previous studies (Kuhl et al., 2014; Rivera-Gaxiola, Klarman, et al., 2005; Rivera-Gaxiola, Silva-Pereyra, et al., 2005) tested the nonnative contrast with the same voiced/pre-voiced distinction but with different consonants (i.e., /d/ vs/ /b/). Results coming from these studies were less consistent. Behaviorally, one study demonstrated 6-8-months-old Spanish-learning infants discriminated the pre-voiced vs. voiced contrast better than English-learning infants (Eilers, Gavin, & Wilson, 1979) while another suggested younger 4-6.5-month Spanish-learning infants could not discriminate this contrast (Lasky, Syrdal-Lasky, & Klein, 1975). Using M/EEG, Rivera-Gaxiola et al. (2005) and the current study observed weaker MMR to this nonnative contrast than the native contrast at 11 months of age while Kuhl et al. (2014) observed the reversed relation. Future research is indeed needed to further elucidate the development of VOT-based speech discrimination.

Some other well-studied nonnative contrasts include the Hindi retroflex/dental contrast, isiZulu bilabial plosive/implosive stop contrast with which both English-speaking adults and older infants have repeatedly shown behaviorally to have extreme difficulty in discriminating (Best & McRoberts, 2003; Werker & Tees, 1984a, 1984b). However, very little is known regarding the neural processes underlying these contrasts. Neural discrimination of a nonnative contrast based on consonant duration was most recently demonstrated to predict later language outcome (Zhao et al., 2021), however, more research with this contrast is needed.

How to conceptually differentiate these nonnative contrasts remains a topic of research and debate. The Perceptual-Assimilation-Model (PAM) combined with the most recent Articulatory-Organ Hypothesis (AOH) proposed by Best provides a framework that emphasizes the articulatory organ involved in producing the nonnative contrast and whether they are used for producing native contrasts (Best, 1994; Best et al., 2016). Another way to conceptualize the differences emphasizes how much acoustic processing/phonetic processing is elicited. It is possible that some nonnative contrasts would index more general auditory processing while others would index more phonetic processing and are more reflective of the NLNC. In the case of the current study, the choice of the pre-voiced vs. voiced bilabial stop contrast may have led to more acoustic processing than phonetic processing.

Future research will need to incorporate multiple nonnative contrasts with various characteristics for a more systematic comparison across them, especially using neuroimaging methods to further disentangle the relative contribution of different neural mechanisms in processing different nonnative contrasts, such as the sensorimotor system and auditory system (Kuhl, 2021). Indeed, the recent MEG studies have all pointed to the left IF region, overlapping with Broca’s area, to be central in speech learning and development, whose function has long been associated with the planning of motor gestures and with linking speech perception and production (Hickok & Poeppel, 2007).

#### 4.4.2 Future comparison to infants with different backgrounds

While understanding the fundamental mechanisms in early speech learning and its relation to language development in typically developing children is foundational to our basic scientific understanding of language acquisition, complementary approaches may also provide invaluable insights. Future research can examine the same developmental trajectories in infants at-risk for developmental communication disorders. By comparing the developmental trajectories between at-risk and typically development infants, we could gain insights into potential early markers of DLDs. Further, future research can also examine the same developmental trajectories in infants who receive music intervention during the sensitive period (e.g. (Zhao & Kuhl, 2016)). A more comprehensive examination will allow us to understand whether such enriched experience can alter the developmental trajectory in ways that may be helpful for infants at-risk for DLD.

## 5. Conclusions

The current study reports on a large-scale longitudinal study on the development of infants’ neural sensitivity to a native and nonnative speech contrast during the ‘sensitive period’ and its relation to later language growth. The results largely replicated and confirmed previous studies showing an enhancement of native contrast discrimination as a group but the nonnative contrast discrimination at 11 months was a better predictor of later individual language outcome. Discrepancies with previous studies were also discussed. Future research is needed to systematically understand infants’ processing of different nonnative speech contrasts and their role in native language acquisition.

## Acknowledgement

This study was supported by a grant from the Bezos Family Foundation. We thank all the research assistants who have devoted years of effort to collect this dataset and note that this work would not be possible without the support of all the participating families. We also thank Dr. Alexis Bosseler for comments on previous versions of the manuscript.

