## Supplemental Table 1 for "Development of infants’ neural speech processing and its relation to later language skills: an MEG study"

Supplemental Material

The 18-month M3L was further excluded from the growth curve analysis due to a flooring effect. Only a marginal trend was observed between 11-month Nonnative MMRs in the left IF and the growth of M3L.

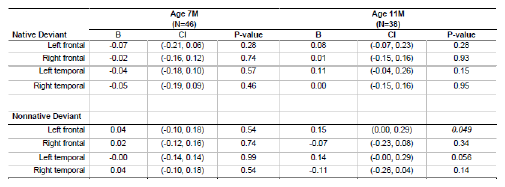

Table S1. MMRs in ROIs predicting growth of M3L from 21 to 30 months of age.
